# Length-weight relationships for 17 fish species in the Luanhe River Estuary, Bohai Sea, northern China

**DOI:** 10.1101/674010

**Authors:** Xu Min, Song Xiao-Jing, Zhang Yun-Ling

## Abstract

Length-weight regressions (LWR, *W* = *a* × *L*^*b*^) of 17 coastal fish species from the Luanhe River Estuary in northern China are presented in this study. A total of 7354 samples from 11 families were measured and weighed. The slope (b) values for LWR ranged from 2.572 in *Acanthogobius ommaturus* to 3.6581 in *Engraulis japonicus*. The median value was 3.114 in *Platycephalus indicus*, although 50% of the values ranged between 2.9451 to 3.2965 for the entire data set.

## Introduction

Estuarine areas are extremely important areas in the life cycle of some fish species. These ecosystems provide food, shelter, and spawning grounds for varieties of marine organisms. The Luan River is a sediment-laden water course on the northern shore of Bohai Bay, China[1]. The estuary of the Luan River is a famous fishing ground and nursery area for marine organisms within Bohai Bay. This area is recognized as an important feeding and breeding location for migratory species[2-5].

Length-weight regressions (LWR) are an important tool for the proper exploitation and management of fish populations[6]. Length and weight data for fish are needed to estimate growth rates, age structure, and other population dynamics[7]. This information is commonly used in the ecosystem modeling approach [8] to calculate the production to biomass ratio (P/B) of different functional groups, taking into account that for more precise weight estimates it is advisable to make use of local values. In addition, LWR allow life history and morphological comparisons between different fish species, or between fish populations from different habitats and/or regions [9]. Biological scientists often estimate fish weight in the field using LWR [10].

Prior to this study there was LWR data available for fish species in the Luanhe River Estuary and this study provides the first LWR references for 17 fish species from this area. This study aimed to provide information that could be used for the management of the Luanhe River fishery grounds. The LWR data will be made available through the Fishbase Database[11], so that they can be used by other researchers.

## Materials and methods

The field survey was approved by the Institute of Oceanology, Chinese Academy of Sciences.

This study was carried out in the Luanhe River Estuary between longitude E 118°57’-119°09’ and latitude N 39°03’-39°15’. The estuary is subject to irregular, semidiurnal tides. The current study was conducted in 2016-2017 as part of a series of studies to assess the biological sustainable capacity of this area. Samples were collected at monthly intervals from December 2016 to August 2017 and at bimonthly intervals from July 2016 to November 2017. The fishing gear used for sampling included a crab pot, trammel net of various inner mesh sizes, and a bottom trawl. Individual trinal nets (50 m in length) with mesh sizes of 2.0 cm, 3.3 cm, 3.5 cm, and 4.0 cm and an outer net mesh of 17.0 cm were used. The trinal nets were 1.3 m high, and the four mesh sizes were connected together, giving a total length of 200 m with four different inner net mesh sizes. The length of a single crab pot was 8 m, and five were connected together at the survey station, giving a total length of 40 m. Trawling was carried out at a speed of 2 knots and the mouth of the net was 2.5 m long.

After hauling, fish samples were immediately transported to the laboratory in Hebei Provincial Research Institute for Engineering Technology of Coastal Ecology Rehabilitation. Specimens were identified to the species level. Scientific names for each species were checked in Fishbase[11]. The standard length (L) of each specimen was measured to the nearest 0.1 cm using a 30-cm ruler. Fish body weight for all specimens was weighed to the nearest 0.01 g using an electric balance (CR-5000WP, Custom, Japan).

LWR were calculated using the equation *W* = *a* × *L*^*b*^ [12]. The relationships between the length and weight of the specimens were calculated by least-square linear regressions applied to logarithmic transformed data combined as[13]:

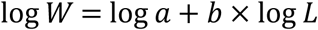

Where ‘W’ is fish body weight (g), ‘L’ is fish standard length (cm), ‘a’ is the initial growth coefficient and ‘b’ is the growth coefficient. The statistical significance level of R^2^ was estimated in LWR fitted by least-squares regression. Only extreme outliers attributed to errors in data collection were omitted from the analyses.

The application of these regressions should be limited to the observed length ranges. These estimated parameters can be treated as mean annual values for the species in our study.

The 95% confidence interval, CI of b was computed using the equation:

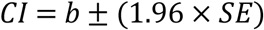

Where *SE* is the standard error of b.

## Results and discussion

A total of 7354 individuals belonging to 17 species (11 families) were recorded in this study. The species, family, sample size (N), length range (cm) and weight range (g), length-weight relationship parameters a and b, 95% confidence interval for b, the coefficient of determination (R^2^) are presented in Table 1. Linear regressions of log transformed data were highly significant (P<0.05) for all analyzed species. The most abundant species sampled was *Chaeturichthys stigmatias* (N=2483). The best represented family was Gobiidae with 4 species recorded.

**Table 1.**
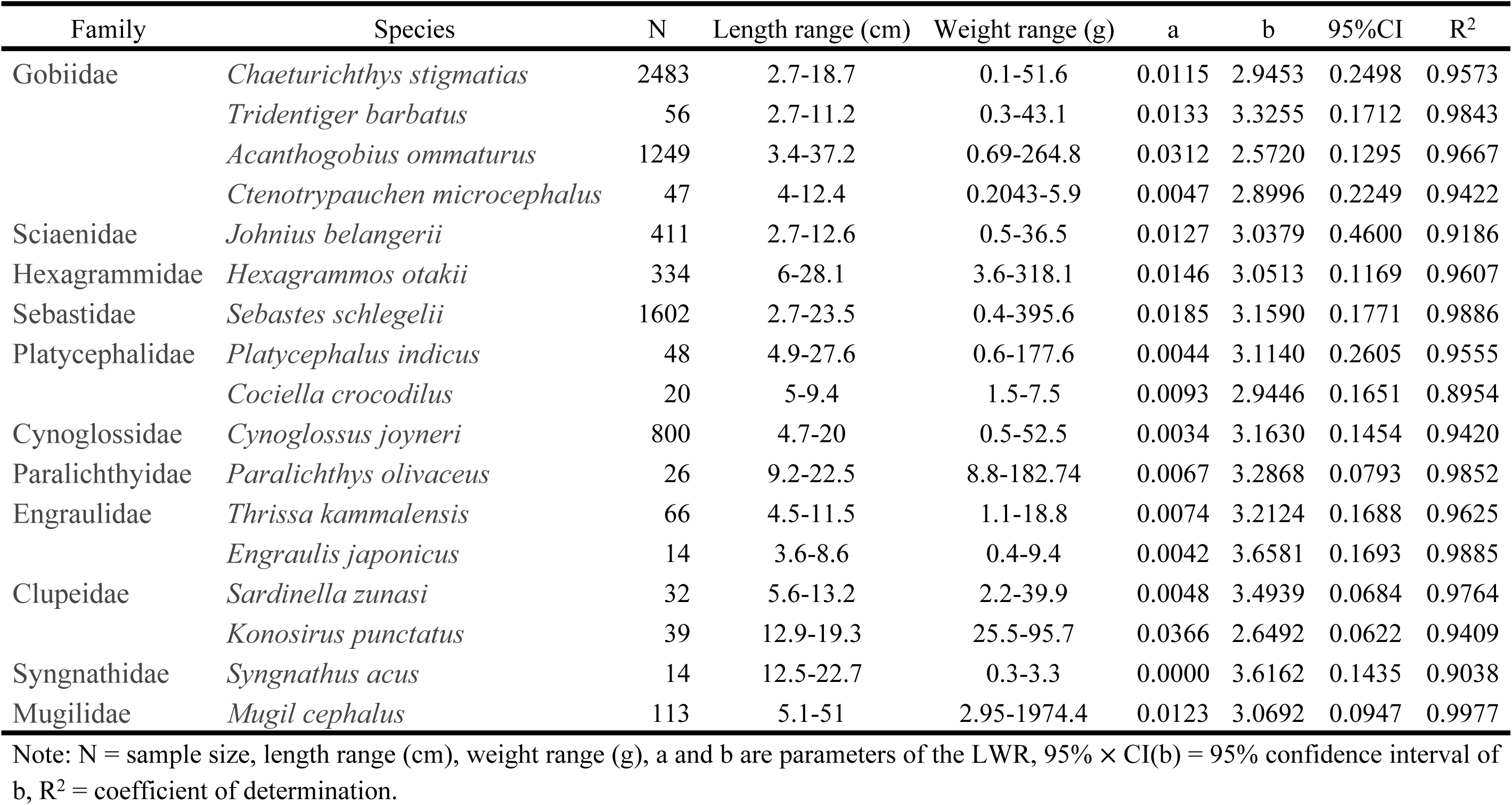
Descriptive statistics and estimated parameters of LWR for 17 fish species caught in the study area.

The coefficients of determination (R^2^) ranged from 0.95 to 1.00 for *Mugil cephalus, Sebastes schlegelii, Engraulis japonicus, Paralichthys olivaceus, Tridentiger barbatus, Sardinella zunasi, Acanthogobius ommaturus, Thrissa kammalensis, Hexagrammos otakii, Chaeturichthys stigmatias, Platycephalus indicus*. While R^2^ values ranged from 0.90 and 0.95 for *Ctenotrypauchen microcephalus, Cynoglossus joyneri, Konosirus punctatus, Johnius belangerii, Syngnathus acus, Cociella crocodilus*, corresponding to a mean value of 0.957(± 0.030).

LWR slope (b) values ranged from 2.572 for *Acanthogobius ommaturus* to 3.6581 for *Engraulis japonicus*. The median value was 3.114 for *Platycephalus indicus*, although 50% of the values ranged from 2.9451 to 3.2965 for the complete data set (Fig.1). When b=3, weight growth is isometric, and when the value of b differs from 3, weight growth is allometric (b>3;b<3). In terms of growth type, these results revealed that 3 species showed negative allometries (b<3), 9 showed positive allometries (b>3) and 5 showed isometric growth (b=3). Most of the species generally presented positive allometric growth. *C. crocodilus* (b=2.9446), *C. stigmatias* (b=2.9453), *J. belangerii* (b=3.0379), *H. otakii* (b=3.0513), *M. cephalus* (b=3.0692) all displayed isometric growth. Various factors may be responsible for differences in parameters of LWR such as temperature, salinity, food (quantity, quality and size), sex, time of year and stage of maturity.

The data collected during this study represents an important contribution of base line data on the LWR of a number of fish species that were previously unavailable. It is important to point out that these LWR should be strictly limited to the length ranges used in the estimation of the linear regression parameters [14]. The results obtained in the current study will contribute to the knowledge of fish populations in the important Luanhe River Estuary and also assist fisheries scientists and managers in the future.

## Acknowledgements

The authors wish to thank Mr Chun Li and the members of Tangshan Marine Ranching L.t.d. for their help with field sampling as well as members from the CAS Key Laboratory of Marine Ecology and Environmental Sciences, Institute of Oceanology, Chinese Academy of Sciences, for constructive discussions and encouragement. The research was supported by Qingdao Postdoctoral Applied Basic Research (No. Y7KY02106N), a Postdoctoral International Exchange Program Introduction Project (No. Y8KY02102L), Zhejiang Provincial Natural Science Foundation of China (Grant No. LY17C190006), and National Natural Science Foundation of China (Grant No. 31702346).

